# Endogenous tagging of Unc-13 reveals nanocluster reorganization at active zones during presynaptic homeostatic potentiation

**DOI:** 10.1101/2022.08.31.506141

**Authors:** Sven Dannhäuser, Achmed Mrestani, Florian Gundelach, Martin Pauli, Fabian Komma, Philip Kollmannsberger, Markus Sauer, Manfred Heckmann, Mila M. Paul

## Abstract

Neurotransmitter release at presynaptic active zones (AZs) requires concerted protein interactions within a dense 3D nano-hemisphere. Among the complex protein meshwork the (M)unc-13 family member Unc-13 of *Drosophila melanogaster* is essential for docking of synaptic vesicles and transmitter release.

We employ MiMIC-based gene editing using GFSTF (EGFP-FlAsH-StrepII-TEV-3xFlag) to endogenously tag all annotated *Drosophila* Unc-13 isoforms enabling visualization of endogenous Unc-13 expression within the central and peripheral nervous system. Electrophysiological characterization using two-electrode voltage clamp (TEVC) reveals that evoked and spontaneous synaptic transmission remain unaffected in *unc-13^GFSTF^* 3^rd^ instar larvae and acute presynaptic homeostatic potentiation (PHP) can be induced at control levels. Furthermore, multi-color structured-illumination shows precise co-localization of Unc-13^GFSTF^, Bruchpilot and GluRIIA-receptor subunits within the synaptic mesoscale. Localization microscopy in combination with HDBSCAN algorithms detect Unc-13^GFSTF^ nanoclusters that move towards the AZ center during PHP with unaltered Unc-13^GFSTF^ protein levels.

## INTRODUCTION

Synapses enable fast and efficient signaling in combination with structural miniaturization (Atwood and Karunanithi, 2002; Chua et al., 2010; Neher and Brose, 2018). Neurotransmitter is released at presynaptic active zones (AZs) consisting of a conserved set of core proteins (Südhof, 2012). One key determinant of synaptic strength is the number of release sites *N* (Silva et al., 2021). The molecular identity of *N* and its dynamics are currently a topic of intense investigations (Neher 2010; Sakamoto et al., 2018; Karlocai et al., 2021). AZs have diameters of 100-1000 nm, thus, super-resolution light microscopy (SLM) is useful to gain insight into their molecular nanotopology. Among the different SLM techniques, localization microscopy allows to quantify protein numbers (Löschberger et al., 2012; Ehmann et al., 2014) and was successfully employed in deciphering AZ nanoarchitecture (Paul et al., 2015; Tang et al., 2016; Sakamoto et al., 2018; Pauli et al., 2021; Mrestani et al., 2021). Most SLM techniques rely on antibodies or, if not available for the epitope of interest, the engineering of genetically tagged fusion proteins. In *Drosophila melanogaster* various tools emerged during the last decade speeding up the generation of genetically marked constructs. Among them is the *Minos-mediated* integration cassette (MiMIC) collection (Venken et al., 2011; Nagarkar-Jaiswal et al., 2015a; Nagarkar-Jaiswal et al., 2015b) and the clustered regularly interspaced short palindromic repeats (CRISPR)-CRISPR-associated protein 9 (Cas9) system (Bassett et al., 2013; Gratz et al., 2013; Yu et al. 2013; Kondo and Ueda, 2013; Gratz et al., 2014; Port et al., 2014). In the MiMIC collection the MiMIC transposon was randomly inserted into the fly genome creating a catalog of over 7000 insertions that facilitated the generation of hundreds of green fluorescent protein (GFP)-tagged constructs (Venken et al., 2011; Nagarkar-Jaiswal et al., 2015a). Some of the insertions are located in AZ genes including *unc-13*. The (M)unc-13 family has homologues in *C. elegans, Drosophila* and mouse (Maruyama and Brenner, 1991; Brose et al., 1995; Aravamudan et al., 1999). Genetic disruption of Munc-13 uncovered an essential role during synaptic release (Richmond et al., 1999; Aravamudan et al., 1999; Augustin et al., 1999). Munc-13 homologues are large multi-domain proteins. Their conserved cysteine-rich C_1_ domain is an internal diacylglycerol and phorbol ester receptor acting in parallel with protein kinase C in the regulation of neurotransmitter secretion (Kazanietz et al., 1995; Betz et al., 1998). Munc-13 interacts with the SNARE-complex protein Syntaxin (Richmond et al., 2001; Madison et al., 2005; Hammarlund et al., 2007) via its C-terminal MUN-domain (Guan et al., 2008). The MUN-domain is surrounded by two C2-domains (C2B and C2C; Liu et al., 2021) and plays an essential role in synaptic transmission including presynaptic homeostatic potentiation (PHP, Böhme et al., 2019; Harrell et al., 2021). Due to its central role in SNARE-complex formation Munc-13 is a key molecular marker of individual release sites (Böhme et al., 2016; Reddy-Alla et al., 2017; Sakamoto et al., 2018; Dittman 2019). Therefore, it would be desirable to visualize all Unc-13 isoforms in identified synapses such as the *Drosophila* neuromuscular junction (NMJ).

We found a promising MiMIC insertion within a coding exon of the fly *unc-13* gene allowing endogenous insertion of a genetic reporter into all predicted variants. We inserted an EGFP-FlAsH-StrepII-TEV-3xFlag (GFSTF) tag into the fly genome using a previously described injection method (Venken et al., 2011). Using electrophysiology, we observed undisturbed neurotransmission and PHP expression in these flies. Furthermore, structured illumination microscopy (SIM) revealed protein trafficking to presynaptic AZs. Finally, combining *direct* stochastic optical reconstruction microscopy (*d*STORM, Heilemann et al., 2008; van de Linde et al., 2011) and hierarchical density-based spatial clustering of applications with noise (HDBSCAN, Campello et al., 2013; Mrestani et al., 2021) we uncover Unc-13^GFSTF^ nanoclusters with ~26 nm diameter at presynaptic AZs that move towards AZ centers during PHP without enhancement of Unc-13^GFSTF^ protein levels, matching the earlier described compaction of other AZ components (Mrestani et al., 2021).

## STAR Methods

### Fly stocks

Flies were raised on standard cornmeal and molasses medium at 25 °C. *Drosophila melanogaster* male 3^rd^ instar larvae of the following strains were used for experiments: Wildtype: *w^1118^* (Bloomington *Drosophila* Stock Center). *unc-13^GFSTF^: y[1] w[*]; Mi{PT-GFSTF.0}unc-13[MI00468-GFSTF.0]*. Original MiMIC-strain (MI00468): *y[1] w[*]; Mi{y[+mDint2]=MIC}unc-13[MI00468]/In(4)ci[D], ci[D] pan[ciD]* (Bloomington *Drosophila* Stock Center #31015).

### Transgene construction

The Minos-mediated integration cassette (MiMIC) strain MI00468 was used to generate the GFP-tagged Unc-13 line (Venken et al., 2011; Nagarkar-Jaiswal et al., 2015a). Microinjection of the plasmid *pBS-KS-attB1-2-PT-SA-SD-0-EGFP-FlAsH-StrepII-TEV-3xFlag (Drosophila* Genomics Resource Center #1298, Venken et al., 2011) and all transgenesis steps were performed at Bestgene Inc. (Chino Hills, USA). Larvae for injection were obtained from crosses of MI00468 males to ΦC31 integrase expressing female virgins. Removal of the genomic ΦC31 integrase source and PCR confirmation of the *attP* sites in MI00468 and of the correct orientation of the RMCE event were performed by the company. *y[1] w[*]; Mi{PT-GFSTF.0}unc-13[MI00468-GFSTF.0] (unc-13^GFSTF^*) was shipped as an unbalanced, homozygous viable stock. Molecular confirmation of precise incorporation of the EGFP-FlAsH-StrepII-TEV-3xFlag (GFSTF, sfGFP used in recent versions of the multi-tag, Venken et al., 2011) tag was performed by genomic PCR with primer pairs *fg_30f* (AATGATAAAGGACAGGGACAAGGT) +*fg_31r* (CTGCTTCATGTGATCGGGGT), *fg50_f* (TGGATGGCGACGTGAAC) +*fg51_r* (GGTTCCATGCAGCATCC) and *fg_58f* (CACAACGTGTACATCACCGC) +*fg_59r* (CTTGAGAACCTGCCGTCCAT). The PCR products were sequenced with *fg_31r, fg50_f* and *fg_58f*.

### Fixation, staining and immunofluorescence

For immunofluorescence imaging of larval neuromuscular junctions and ventral nerve cords, larvae were dissected in ice-cold hemolymph-like solution (HL-3, Stewart et al., 1994), fixed with 4 % paraformaldehyde (PFA) in phosphate buffered saline (PBS) for 10 minutes and blocked for 30 minutes with PBT (PBS containing 0.05 % Triton X-100, Sigma) including 5 % natural goat serum (NGS, Dianova). Primary antibodies were added for overnight staining at 4 °C. After two short and three long washing steps with PBT (60 min each for data shown in Figure 2C, 20 min each for all other imaging data), preparations were incubated with secondary antibodies for 3 hours at room temperature, followed by two short and three 20 min long washing steps with PBT. Preparations were kept in PBS at 4 °C until imaging. All data were obtained from NMJs formed on abdominal muscles 6/7 in segments A2 and A3. Directly compared data were obtained from larvae stained in the same vial and measured in one imaging session. Primary antibodies were used in the following concentrations: mouse α-Brp (Brp^Nc82^, 1:100; AB_2314866, Developmental Studies Hybridoma Bank), rabbit-α-GFP (1:1,000 for *d*STORM or 1:3,000 for Zeiss Axiovert and structured illumination microscopy; A11122, Thermofisher), mouse α-GluRIIA (1:100; AB_528269, Developmental Studies Hybridoma Bank) and Cy5-conjugated α-horseradish-peroxidase (α-HRP, 1:250, AB_2338714, Jackson ImmunoResearch).

**Figure 1.**
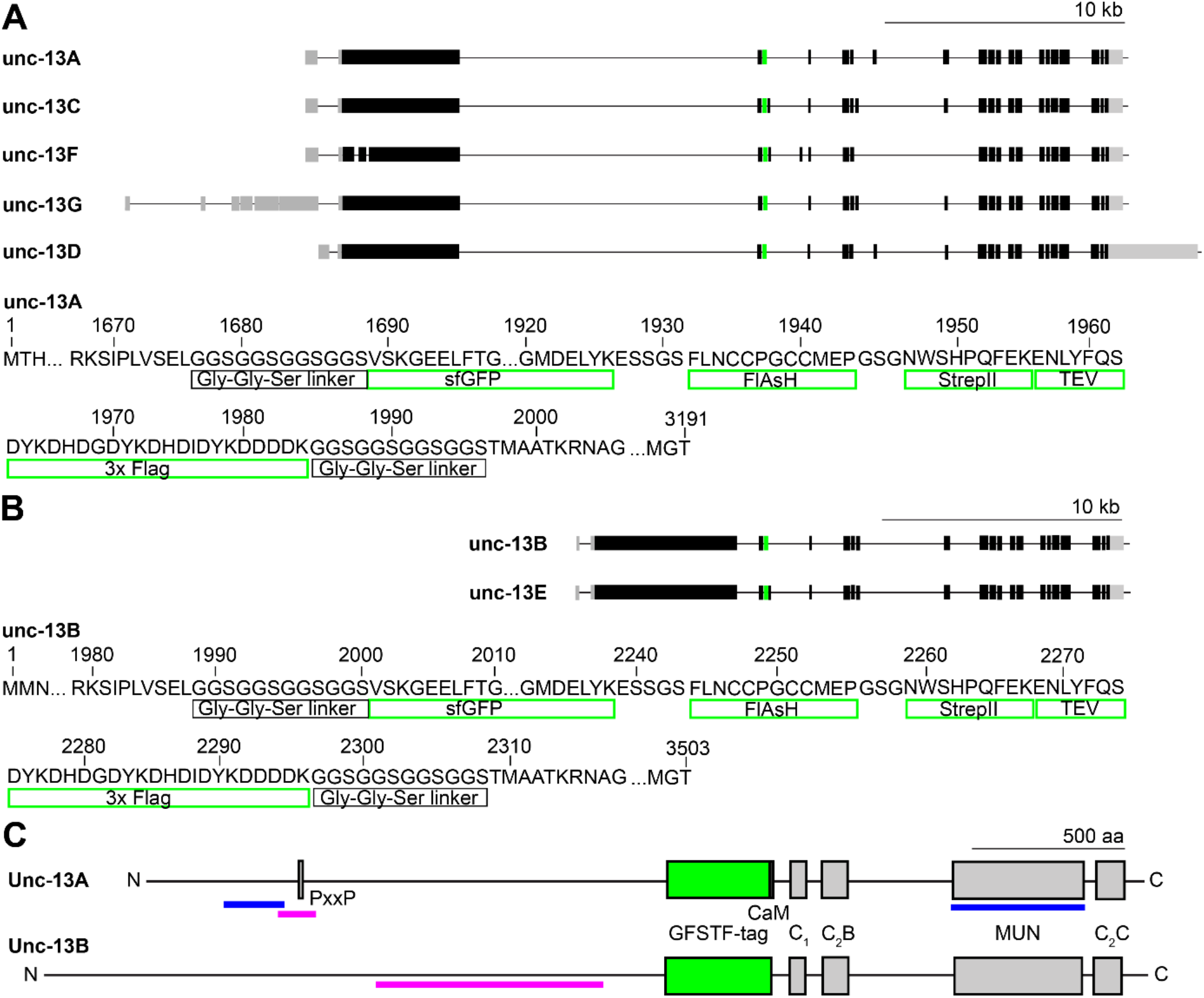
Endogenous tagging of *unc-13* in *Drosophila melanogaster*. (A, B) Genetic map of *unc-13* in *Drosophila* with the 5 annotated transcripts A, C, F, G and D (A) and the 2 annotated transcripts B and E (B). Black boxes indicate exons, grey boxes introns and green boxes mark the insertion site of the GFSTF-tag located in a coding intron. Lower panels show amino acids of the Unc-13A and Unc-13B fly protein illustrating the specific elements the GFSTF-tag consists of. (C) Schematic illustration of Unc-13A and Unc-13B domains: CaM- (Calmodulin), C_1_-, C_2_B-, C_2_C- and MUN-domain and the PxxP-motif. Green box marks GFSTF-tag, magenta bars indicate epitopes of Unc-13A and Unc-13B antibodies (Böhme et al., 2016) and blue bars of N- and C-term antibodies, respectively (Reddy-Alla et al., 2017).

**Figure 2.**
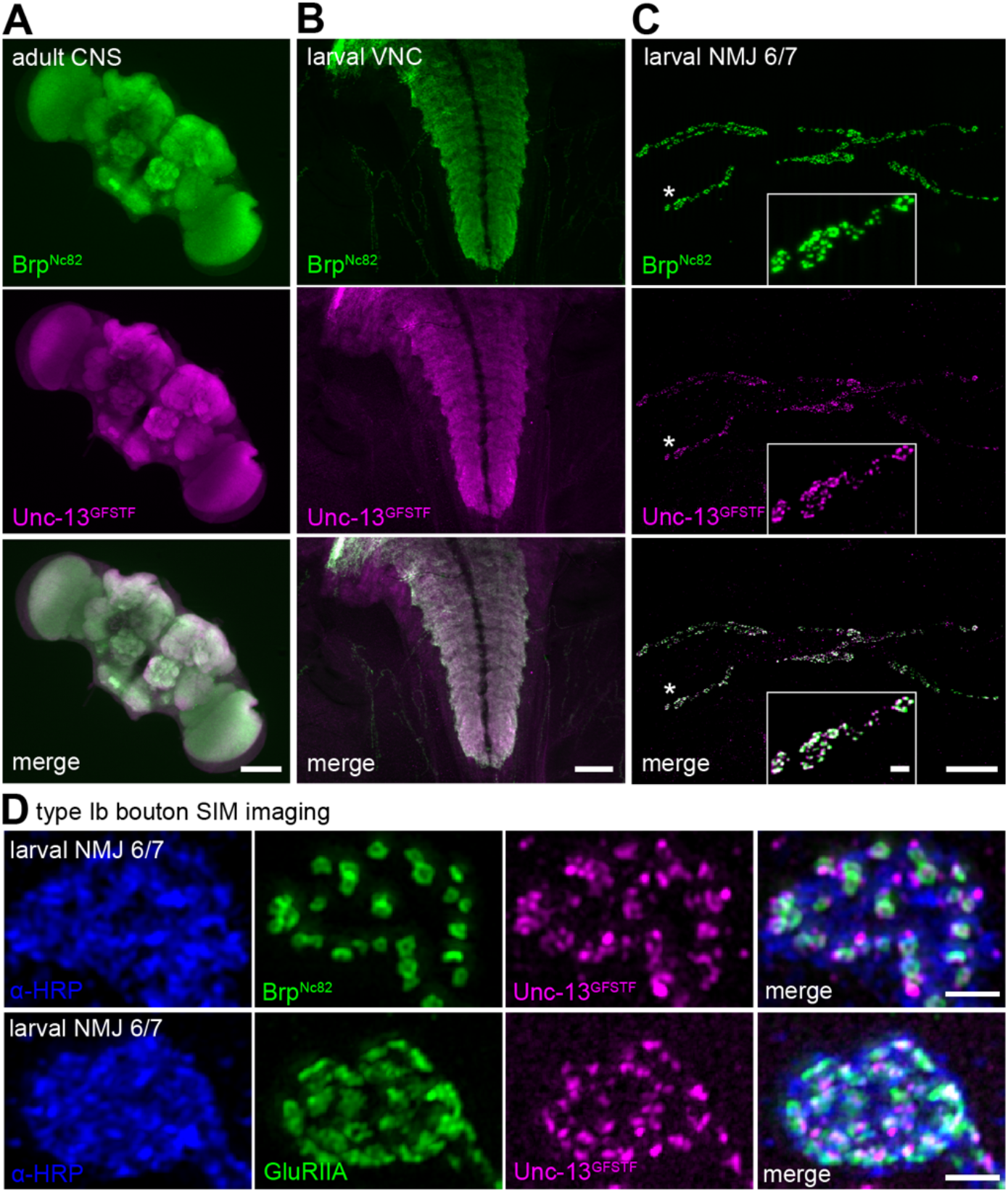
Unc-13^GFSTF^ is expressed in the adult and larval central nervous system and at larval NMJs. (A) Adult central nervous system (CNS) stained with Brp^Nc82^ (green) and rabbit α-GFP antibody to visualize Unc-13^GFSTF^ (magenta). (B) Larval ventral nerve cord (VNC) of a male 3^rd^ instar *Drosophila* larva stained with the same antibodies as in (A). (C) Co-expression of Brp^Nc82^ (green) and Unc-13^GFSTF^ (magenta) at a 3^rd^ instar larval neuromuscular junction (NMJ) on abdominal muscles 6/7 in segment A3. (D) Structured illumination microscopy imaging of a type Ib bouton of a larval NMJ on abdominal muscles 6/7 shows pre- and postsynaptic co-localization of α-HRP (blue), Unc-13^GFSTF^ (magenta) and Brp^Nc82^ (green, upper panel) or GluRIIA receptor subunits (green, lower panel). Scale bars in (A) and (B) 100 μm, in (C) 20 μm and 2 μm (inset) and in (D) 1 μm.

For immunofluorescence imaging of adult brains, fly heads were dissected and brains removed in ice-cold PBS, fixed in 4 % PFA in PBS for 30 minutes and blocked for 30 minutes with 0.6 % PBT including 5 % NGS. Incubation in primary antibodies, washing in 0.6 % PBT, incubation in secondary antibodies and again washing followed the protocol explained in detail above. The same primary antibodies were used in the following concentrations: mouse α-Brp (Brp^Nc82^, 1:100) and rabbit-α-GFP (1:1,000).

### Axiovert imaging and structured illumination microscopy (SIM)

Preparation, fixation and staining was performed as described above. Secondary antibodies were used in the following concentrations for larval neuromuscular junctions, ventral nerve cords and adult fly brains: goat α-rabbit conjugated Cy3 (1:500, AB_2338006, Jackson ImmunoResearch) and goat α-mouse conjugated Alexa Fluor488 (1:500, A32723, Invitrogen). Larval preparations were mounted in Vectashield (Vector Laboratories, Burlingame, CA) for Axiovert imaging or Prolong Glass (ThermoFisher) for structured illumination microscopy. Images were acquired at room temperature from NMJs on muscles 6/7 in segments A2 and A3. An Apotome System (Zeiss, Axiovert 200M Zeiss, objective 63x, NA 1.4, oil) was used for low-resolution characterization of larval NMJs, larval ventral nerve cords and adult brains (Figure 2A-C). For SIM we used a Zeiss Elyra S.1 structured illumination microscope equipped with a sCMOS camera (pco.edge 5.5 m) and an oil-immersion objective (Plan-Apochromat 63x, 1.4 NA). Lasers with 488 nm, 531 nm and 641 nm wavelength were used. Z step size was set to 0.1 μm, and imaging was performed using five rotations of the grating at five different phase steps. Fourier transformation of structured illumination images was performed using ZEN software (Carl Zeiss, Jena), and subsequent analysis was done with ImageJ.

### *direct* stochastic optical reconstruction microscopy (*d*STORM)

*d*STORM imaging of the specimen was performed essentially as previously reported (Ehmann et al., 2014; Paul et al., 2015; Mrestani et al., 2021; Paul et al., 2022). Preparations were incubated with respective primary antibodies as described above. The following secondary antibodies were used: goat α-rabbit F(ab’)2 fragments labelled with Alexa Fluor647 (1:500; A21246, Thermofisher) and goat α-mouse IgGs labelled with Alexa Fluor532 (1:500; A11002, Thermofisher). After staining, larval preparations were incubated in 100 mM mercaptoethylamine (MEA, Sigma-Aldrich) in a 0.2 M sodium phosphate buffer, pH ~7.9, to allow reversible switching of single fluorophores during data acquisition (van de Linde et al., 2008). Images were acquired using an inverted microscope (Olympus IX-71, 60x, NA 1.49, oil immersion) equipped with a nosepiece-stage (IX2-NPS, Olympus). 647 nm (F-04306-113, MBP Communications Inc.) and 532 nm (gem 532, Laser Quantum) lasers were used for excitation of Alexa Fluor647 and Alexa Fluor532, respectively. Laser beams were passed through clean-up filters (BrightLine HC 642/10 and Semrock, ZET 532/10, respectively), combined by two dichroic mirrors (LaserMUX BS 514-543 and LaserMUX BS 473-491R, 1064R, F38-M03, AHF Analysentechnik), and directed onto the probe by an excitation dichroic mirror (HC Quadband BS R405/488/532/635, F73-832, AHF Analysentechnik). The emitted fluorescence was filtered with a quadband-filter (HC-quadband 446/523/600/677, Semrock) and a long pass- (Edge Basic 635, Semrock) or bandpass-filter (Brightline HC 582/75, Semrock) for the red and green channel, respectively, and divided onto two cameras (iXon Ultra DU-897-U, Andor) using a dichroic mirror (HC-BS 640 imaging, Semrock). For the red channel, image resolution was 127 nm x 127 nm per pixel to obtain super-resolution of Unc-13^GFSTF^. For the green channel, image resolution was 130 nm x 130 nm per pixel. Single fluorophores were localized and high resolution-images were reconstructed with rapi*d*STORM (Heilemann et al., 2008; van de Linde et al., 2011; Wolter et al., 2010; Wolter et al., 2012; www.super-resolution.de). Only fluorescence spots with an A/D count over 12,000 were analyzed and a subpixel binning of 10 nm px^-1^ was applied.

### Analysis of localization data

Localization data were analyzed essentially as described previously (Mrestani et al., 2021) with an extension for two-channel localization data. Analysis was performed with custom written Python code (https://www.python.org/, language version 3.6) and the web-based Python interface Jupyter (https://jupyter.org/index.html). Localization tables from rapi*d*STORM were directly loaded and analyzed. Prior to the Python-based analysis the regions of interest (ROI) were masked in the reconstructed, binned images from rapi*d*STORM using FIJI (Schindelin et al., 2012). These ROIs corresponded to the terminal 6 boutons. For cluster analysis we used the Python implementation of HDBSCAN (McInnes et al., 2017; https://github.com/scikit-learn-contrib/hdbscan) which takes ‘minimum cluster size’ and ‘minimum samples’ as the main free parameters. In the first step we extracted Brp clusters in the Alexa Fluor532 channel, corresponding to individual AZs, with the combination 100 and 25 for minimum cluster size and minimum samples (Figure 4Aiii, compare Mrestani et al., 2021). All unclustered localizations were discarded from further analysis. Brp clusters were used to remove noise from the Unc-13^GFSTF^ Alexa Fluor647 channel such that all Unc-13^GFSTF^ localizations with an euclidian distance > 20 nm to Brp localizations were discarded. The H function (Figure 4B) as derivative of Ripley’s K function was computed using Python package Astropy (Robitaille et al., 2013) according to our previous algorithm (Mrestani et al., 2021) for the denoised Unc-13^GFSTF^ localizations and for the random Poisson distribution. Curves for display were averaged (mean ± SD). The function was evaluated in nm steps for radii from 0 to 120 nm and without correction for edge effects. A second HDBSCAN to extract the individual Unc-13^GFSTF^ subclusters (SC) was performed with minimum cluster size 7 and minimum samples 2 to get SCs with a radius that matches the maximum of the H function. Unc-13^GFSTF^ SCs were assigned to individual Brp clusters by computing the euclidian distance between the center of mass (c.o.m.) of each SC and the c.o.m. of each Brp cluster and selecting the lowest distance. In that way, the number of SCs per AZ could be quantified. The distance between the c.o.m.s is referred to as radial distance and was computed for each individual AZ as mean of the assigned Unc-13^GFSTF^ SCs. To quantify cluster areas, we computed 2D alpha shapes using CGAL (Computational Geometry Algorithms Library, https://www.cgal.org) in Python. To get the alpha shapes of Brp clusters and Unc-13^GFSTF^ SCs we choose α-values of 800 nm^2^ and 300 nm^2^, respectively. Exclusion criteria for Brp clusters were area < 0.03 μm^2^ and > 0.3 μm^2^ (Mrestani et al., 2021). Unc-13^GFSTF^ SCs that were assigned to those Brp clusters were also excluded from further analysis, as well as SCs where alpha shape determination failed due to sparse signal that yielded SC areas of 0 μm^2^. The Unc-13 area per AZ (Figure 4E) was computed as the sum of all Unc-13 SC areas belonging to an individual AZ. Brp cluster circularity was computed as described previously (Mrestani et al., 2021). For the analysis of Unc-13^GFSTF^ superclusters (SpC, Figure 5B and E-H) and distances between SC c.o.m.s (Figure 5C and D) only AZs with a circularity ≥ 0.6 were selected. The Python module scikit-learn (Pedregosa et al., 2011) was used to compute distances between SC c.o.m.s. To extract SpCs, HDBSCAN was performed taking the SC c.o.m.s of an individual AZ as input (minimum cluster size and minimum samples 2, respectively). The cluster selection method was changed to ‘leaf’ clustering, which comprises a tendency to more homogeneous clusters by extracting those that lie on leaf nodes of the cluster tree rather than the most stable clusters. The default setting ‘excess of mass’, that was used for all other cluster analyses throughout this study, delivered similar statistical results and median values (data not shown) but less intuitive clustering. For the statistical comparison of SpC numbers per AZ between experimental groups (Figure 5F) only AZs where SpCs could be detected were included. The SpC c.o.m. was defined as the c.o.m. of its respective SC c.o.m.s and the euclidian distance between these SC c.o.m.s and the SpC c.o.m. was computed as mean per SpC.

**Figure 3.**
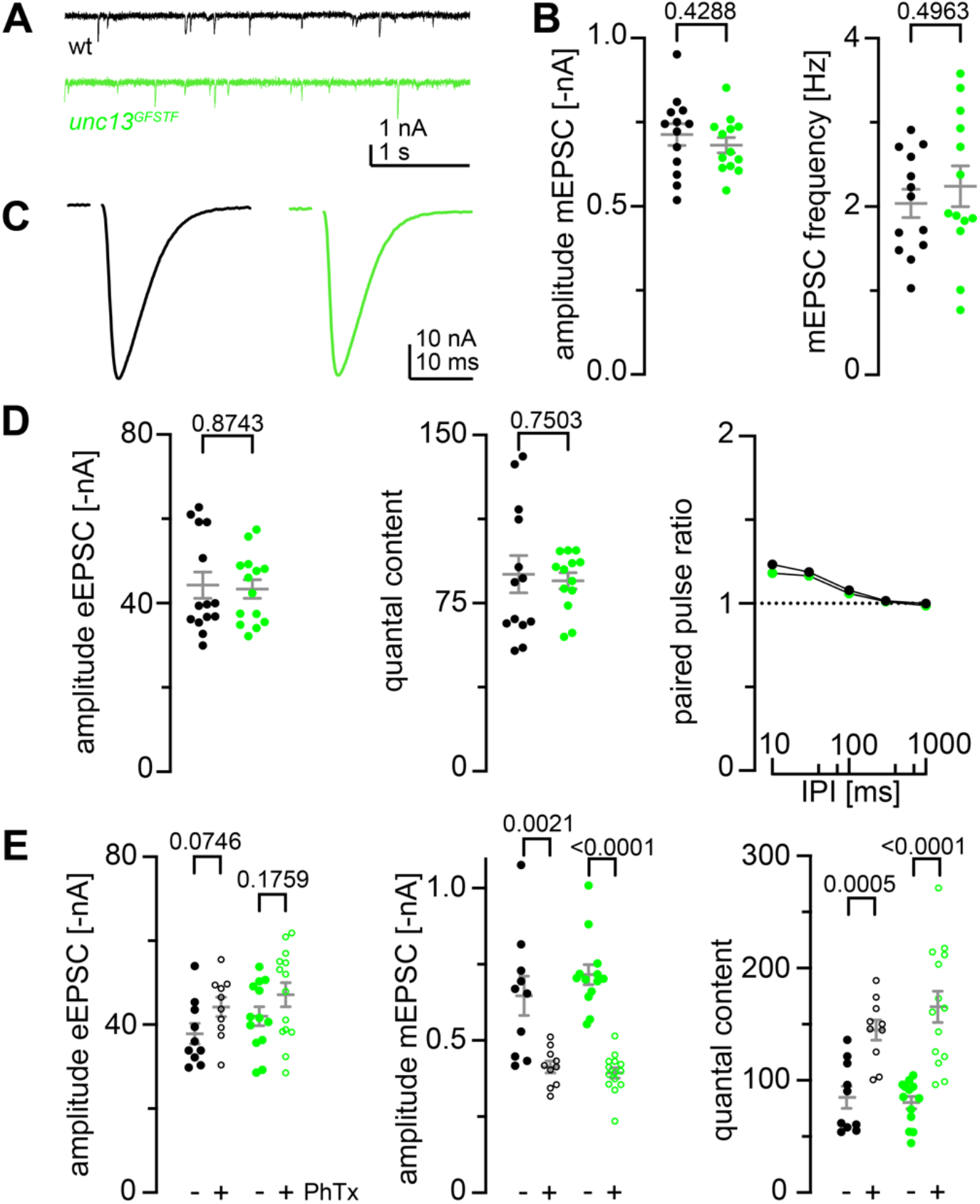
Endogenous tagging of *unc-13* leaves synaptic function and presynaptic homeostatic potentiation undisturbed. (A) Representative traces of miniature EPSCs recorded in 1 mM Ca^2+^ at wildtype (wt, black) and *unc-13^GFSTF^* (green) NMJs. (B) mEPSC amplitude and frequency are unaltered in *unc-13^GFSTF^* larvae (green, n = 13 NMJs from 7 larvae) compared to wt (black, n = 13 NMJs from 8 larvae) (C) Representative traces of evoked EPSCs recorded in 1 mM extracellular Ca^2+^ in both genotypes. (D) eEPSC amplitude, quantal content and paired-pulse-ratios measured with different interpulse intervals (IPI; 10, 30, 100, 300 and 1000 ms) remain unaltered in *unc-13^GFSTF^* animals (wt: n = 14 NMJs from 8 larvae; *unc-13^GFSTF^:* n = 14 NMJs from 7 larvae). (E) eEPSC amplitude, mEPSC amplitude and quantal content in wt (black) and *unc-13^GFSTF^* (green) animals treated with PhTx in dmso (+, open circles) or dmso (-, filled circles). *unc-13^GFSTF^* larvae still exhibit presynaptic homeostatic potentiation in response to PhTx stimulation (wt: 10 NMJs from 7 larvae in dmso, 10 NMJs from 5 larvae in PhTx; *unc-13^GFSTF^*: 13 NMJs from 6 larvae in dmso, 14 NMJs from 7 larvae in PhTx). Whisker plots represent mean ± SEM, scatter plots show individual data points, individual p-values are indicated.

**Figure 4.**
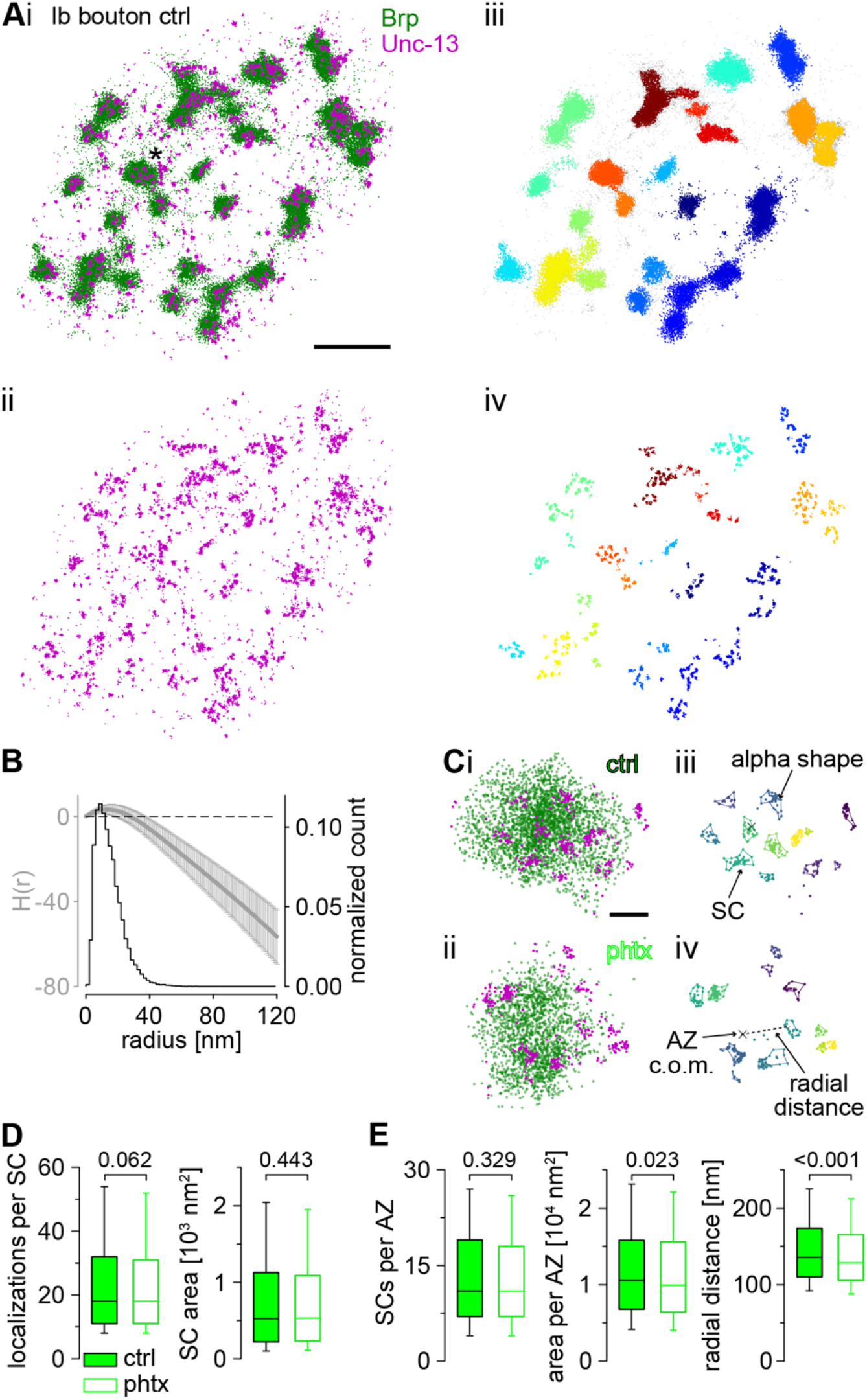
Super-resolution imaging of Unc-13^GFSTF^ at presynaptic AZs reveals nanoscale reorganization during PHP. (A) Scatter plots of two-channel *d*STORM localizations of an *unc-13^GFSTF^* type Ib bouton from a control (ctrl) animal co-stained with α-GFP antibody labelled with Alexa Fluor647 conjugated F(ab’)2 fragments for visualization of Unc-13^GFSTF^ (magenta) and Brp^Nc82^ labelled with Alexa Fluor532 conjugated IgGs (green). (Ai) Overlay of the two channels. Asterisk marks the enlarged region in (Ci). (Aii) Red channel showing Unc-13^GFSTF^ localizations of the same bouton as in (Ai). (Aiii) Brp localizations from (Ai) with clusters extracted by HDBSCAN and individual AZs in different colors. Unclustered localizations are shown in grey. (Aiv) Unc-13^GFSTF^ localizations from (Aii) with all localizations with euclidian distance > 20 nm to Brp localizations removed. The removed signal is considered noise. Individual Unc-13^GFSTF^ subclusters (SCs) were extracted by HDBSCAN and assigned to nearest AZs by color. (B) Averaged H function (black, mean ± SD) from n = 2,040 Unc-13^GFSTF^ first level clusters obtained from 22 NMJs from 9 animals (maximum of the curve indicates a mean sc. radius of 13 nm) and histogram (grey) of the mean radius from n = 20,037 Unc-13^GFSTF^ SCs (estimated from SC area under the assumption of a circular area, median (25^th^-75^th^ percentile): 12.9 (8.4-18.9) nm). Dashed black line indicates the prediction for a random Poisson distribution. (C) Scatter plots of Ib AZs from a ctrl (Ci, iii) and a Philanthotoxin treated animal (phtx, Cii, iv). (Ci, ii) Original scatter plots Unc-13^GFSTF^ and Brp^Nc82^ *d*STORM localizations. Unclustered localizations are not shown in both channels. (Ciii, iv) Unc-13^GFSTF^ SCs extracted by HDBSCAN, that are assigned to an individual ctrl (Ciii) and phtx (Civ) AZ, in different colors. Colored lines indicate alpha shapes used for area determination. Grey dots are unclustered localizations. Centers of mass (c.o.m.) of the corresponding AZ (cross) are indicated. Dashed line shows the euclidian distance between the AZ c.o.m and an SC c.o.m., referred to as radial distance. (D) Number of localizations per Unc-13^GFSTF^ SC and SC area in ctrl (filled boxes, n = 20,037 SCs from 22 NMJs and 9 animals) and phtx (open boxes, n = 20,393 SCs from 23 NMJs and 9 animals) shown as box plots (horizontal lines show median, box boundaries 25^th^ and 75^th^ percentiles, whiskers 10^th^ and 90^th^ percentiles). (E) Number of Unc-13^GFSTF^ SCs, total Unc-13^GFSTF^ area and radial distance of Unc-13^GFSTF^ SCs per AZ in ctrl (n = 1,462 AZs) and phtx (n = 1,521 AZs). Scale bars in (A) 1 μm, in (C) 100 nm.

**Figure 5.**
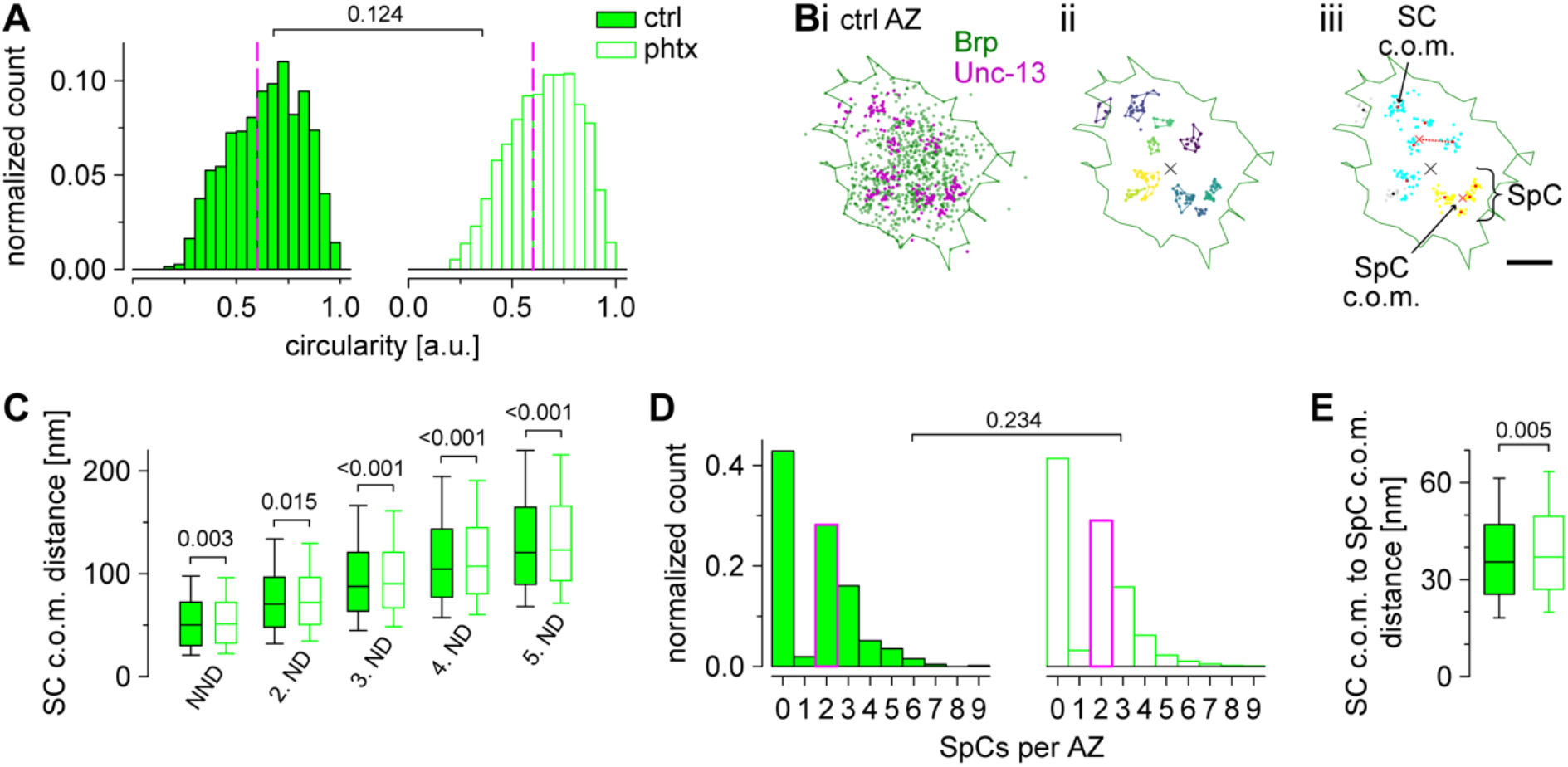
Unc-13^GFSTF^ superclusters are enlarged during PHP. (A) AZ circularity computed for Brp clusters in *unc-13^GFSTF^* after dmso (control, ctrl; filled bars, n = 1,462 AZs from 22 NMJs and 9 animals) or PhTx treatment (open bars, n = 1,521 AZs from 23 NMJs and 9 animals). Dashed magenta lines indicate the cutoff circularity of ≥ 0.6 a.u. used for further analyses. (B) Scatter plots of a representative, circular AZ from a ctrl type Ib bouton. Green line indicates the alpha shape used for area determination of the AZ. (Bi) Original scatter plots of Unc-13^GFSTF^ and Brp^Nc82^ *d*STORM localizations. (Bii) Unc-13^GFSTF^ SCs extracted by HDBSCAN in different colors. Unclustered localizations are not shown. Colored lines indicate distinct alpha shapes. The center of mass (c.o.m.) of the corresponding AZ (black cross) is shown. (Biii) Two individual superclusters (SpCs), indicated by blue and yellow colors, extracted by HDBSCAN from the c.o.m.s (red dots) of Unc-13^GFSTF^ SCs shown in (Bii). Black dots represent SC c.o.m.s that are unclustered according to HDBSCAN analysis and grey dots show localizations of the corresponding SCs. Red crosses indicate c.o.m.s of SpCs. Dashed red line indicates the distance between a SC c.o.m. and its respective SpC c.o.m. (C) Nearest neighbor distance (NND) and 2^nd^ to 5^th^ neighbor distance (ND) of Unc-13^GFSTF^ SCs at circular AZs for ctrl (n = 10,461, 10,425, 10,329, 10,185 and 9,945 distances from 876, 858, 826, 790 and 742 AZs for NND and 2^nd^ to 5^th^ ND, respectively) and phtx (n = 10,988, 10,940, 10,856, 10,688 and 10,493 distances from 927, 903, 875, 833 and 794 AZs for NND and 2^nd^ to 5^th^ ND, respectively). (D) Number of SpCs per AZ. Median values, indicated in magenta, as well as statistical comparison refers to AZs with at least one SpC (ctrl: n = 501 from 22 NMJs and 9 animals; phtx: n = 543 AZs from 23 NMJs and 9 animals). (E) Distance between c.o.m.s of SCs and SpCs for both groups at AZs with at least one SpC (n = 1,417 and 1,497 SpCs for ctrl and phtx, respectively).

### Electrophysiology

TEVC recordings (Axoclamp 2B amplifier, Digidata 1440A; Molecular Devices) were obtained from abdominal muscle 6 in segments A2 and A3 as previously described (Paul et al., 2022). All measurements were obtained at room temperature in HL-3 (Stewart et al., 1994) with the following composition (in mM): NaCl 70, KCl 5, MgCl_2_ 20, NHCO^3^ 10, trehalose 5, sucrose 115, Hepes 5, and CaCl_2_ 1, pH adjusted to 7.2. Intracellular electrodes had resistances of 10–30 MΩ and were filled with 3 M KCl. For analysis, only cells with an initial membrane potential of at least −50 mV and a membrane resistance of ≥4 MΩ were included. During recordings, cells were clamped at a holding potential of −80 mV (minis) or −60 mV (evoked EPSCs). To evoke synaptic currents, nerves were stimulated via a suction electrode with pulses of 300 μs length and typically at 12 V (Grass S48 stimulator and isolation unit SIU5; Astro-Med). Signals were low-pass filtered at 10 kHz and analyzed in Clampfit 11.1 (Molecular Devices). Paired-pulse recordings were performed with interstimulus intervals of (in ms: 10, 30, 100, 300, and 1,000). Between recordings, cells were given a 10 s rest. For analysis, 5 - 10 traces per interval were averaged. The amplitude of the second response in 10 ms interpulse recordings was measured from the peak to the point of interception with the extrapolated first eEPSC as described previously (Hallermann et al., 2010; Weyhersmüller et al., 2011). To assess basal synaptic transmission 10 EPSCs evoked at 0.2 Hz were averaged per cell. The quantal content was estimated by dividing the mean eEPSC amplitude by the mean mEPSC amplitude measured in the same cell. mEPSC amplitudes were corrected for the more hyperpolarized holding potential (Hallermann et al., 2010).

### Philanthotoxin treatment

Philanthotoxin 433 tris (trifluoroacetate) salt (PhTx, CAS 276684-27-6, Santa Cruz Biotechnology) was dissolved in dimethyl sulfoxide (DMSO) to obtain a stock solution of 4 mM and stored at −20 °C. For each experiment, the respective volume was further diluted with freshly prepared HL-3 to a final PhTx concentration of 20 μM in 0.5 % DMSO. Control experiments were performed with the same DMSO concentration in HL-3. PhTx treatment of semi-intact preparations was performed essentially as described previously (Frank et al., 2006; Mrestani et al., 2021). In brief, larvae were pinned down in calcium-free, ice-cold HL-3 at the anterior and posterior endings, followed by a dorsal incision along the longitudinal axis. Larvae were incubated in 10 μl of 20 μM PhTx in DMSO for 10 minutes at room temperature. Following this incubation time, PhTx was replaced by HL-3 and larval preparations were completed, followed by electrophysiological measurements or *d*STORM imaging.

### Statistics

Statistical analyses were performed with Sigma Plot 13 (Systat Software) or GraphPad Prism 9 (San Diego, CA). Shapiro-Wilk was used to test normality. If data were not normally distributed, we used the non-parametric Mann-Whitney rank sum test for statistical analysis and reported data as median (25^th^-75^th^ percentile). If data were normally distributed, they were reported as mean ± SD unless indicated otherwise. In box plots, horizontal lines represent median, boxes quartiles and whiskers 10^th^ and 90^th^ percentiles. Scatter plots show individual data points unless indicated otherwise.

Bin counts in histograms were normalized to the total number of observed events which was set to 1. All plots were produced with Sigma Plot. Figures were assembled using Adobe Illustrator (Adobe, 2015.1.1 release). Supplementary Tables 1–4 contain all numerical values not stated in text and figure legends including p-values and samples sizes.

### Code and data availability

The authors declare that custom written Python code and all data sets supporting the findings of this work are available from the corresponding authors.

## RESULTS

### Endogenous tagging of unc-13

To generate an endogenously tagged *unc-13* locus, we screened the previously established MiMIC library (Venken et al., 2011; Nagarkar-Jaiswal et al., 2015a; Nagarkar-Jaiswal et al., 2015b) for an appropriate line. The MiMIC transposon harbors selection markers (*yellow*^+^ and enhanced GFP [EGFP]) and two inverted *attP* sites for recombinase-mediated cassette exchange (RMCE), as well as a gene-trap cassette (Venken et al., 2011). The insertion MI00468 carries a MiMIC element in a coding intron and, via RMCE with a protein-trap plasmid, allows the generation of a new exon that gets spliced in all annotated *unc-13* transcripts (Figure 1A, B). This strategy was applied to insert a multi-tag, consisting of superfolder (sf) GFP (sfGFP), a modified Fluorescein arsenical helix binder (FlAsH) binding tetracysteine motif, the StrepII peptide tag, a TEV protease cleavage site and the Flag peptide tag (EGFP-FlAsH-StrepII-TEV-3xFlag [GFSTF] tag), using a previously published protein-trap plasmid (see Material and Methods; Venken et al., 2011; Nagarkar-Jaiswal et al., 2015). The multi-tag combines the advantages of different protein and peptide tags including several possibilities for antibody staining, live imaging and RNAi knockdown. GFSTF is inserted at the same amino acid positions in the two major isoforms Unc-13A (Figure 1A) and Unc-13B (Figure 1B) but is C-terminally followed by alternative amino acids in Unc-13C, E and F, resulting from the additional exon in the respective transcripts. The multi-tag is located at the C-terminal side of the large unstructured N-Terminus and is followed by all annotated domains of the proteins (Figure 1C). The animals are homozygous viable and appear healthy (e.g., emerging and effective flight). In the following, we focus on the detection of sfGFP using a combination of primary IgG antibodies and secondary F(ab’)_2_ fragments to establish a protocol comparable to earlier applications of *d*STORM imaging at the *Drosophila* NMJ (Ehmann et al., 2014; Paul et al., 2015; Mrestani et al., 2021; Paul et al., 2022). We refer to the fusion protein and the genotype as Unc-13^GFSTF^ and *unc-13^GFSTF^*, respectively.

### Expression of Unc-13^GFSTF^ in the adult and developing *Drosophila* nervous system

To assess expression of the endogenously tagged Unc-13 in the *Drosophila* nervous system, we performed immunostainings using a polyclonal antibody against GFP (see Material and Methods) for detection of the sfGFP within the GFSTF-tag and a well characterized, highly specific monoclonal antibody Brp^Nc82^ mapping to the C-terminal region of Bruchpilot (Brp, Kittel et al., 2006; Fouquet et al., 2009, Figure 2). We found strong and specific expression of Unc-13^GFSTF^ in the adult and larval central nervous system (Figure 2A, B). In addition, we detected reliable Unc-13^GFSTF^ expression at larval NMJs in co-expression with Brp (Figure 2C). Since Brp is an abundant presynaptic epitope and covers the spatial extent of an individual presynaptic AZ (Mrestani et al., 2021) we set out to analyze the co-localization of Brp and Unc-13^GFSTF^. We employed structured illumination microscopy (SIM) at individual type Ib boutons. Our data revealed co-localization of Unc-13^GFSTF^ and Brp at presynaptic AZs (Figure 2D) and of Unc-13^GFSTF^ and postsynaptic glutamate receptors using a polyclonal antibody against the GluRIIA subunit (Figure 2E). We conclude that incorporation of the fluorescent GFSTF-tag at the endogenous locus of *unc-13* in *Drosophila* reports reliable and reasonable localization of the protein at presynaptic AZs allowing further investigations at the nanoscale level (compare Figures 4 and 5).

### Normal synaptic function of *unc-13^GFSTF^* NMJs

To evaluate spontaneous and evoked synaptic release at larval NMJs of *unc-13^GFSTF^* we performed two-electrode voltage clamp recordings (TEVC, Figure 3). First, miniature excitatory postsynaptic currents (mEPSCs) were recorded to examine spontaneous vesicle release (Figure 3A). mEPSC amplitudes and the frequency of spontaneous fusion events were unchanged in *unc-13^GFSTF^* compared to wildtype controls (Figure 3B). In addition, evoked excitatory postsynaptic currents (eEPSCs) in response to nerve stimulation were unaltered in *unc-13^GFSTF^* compared to controls (Figure 3C, D). Furthermore, quantal content was unchanged in *unc-13^GFSTF^* (Figure 3D). Next, we tested whether insertion of the tag alters synaptic short-term plasticity. We found paired pulse ratios (PPR) unchanged in *unc-13^GFSTF^* (Figure 3D). Taken together, these data reveal no differences between *unc-13^GFSTF^* and control larvae in basal transmission properties of spontaneous and evoked synaptic release, making the endogenously tagged Unc-13 variant a valuable tool for the assessment of AZ function and structure.

### Presynaptic homeostatic potentiation in *unc-13^GFSTF^*

In addition to its role in docking and priming of synaptic vesicles Unc-13 is involved in diverse presynaptic plasticity processes including Ca^2+^, DAG or RIM-dependent short-term plasticity (Betz et al., 2001; Dulubova et al., 2005; Shin et al., 2010; Xu et al., 2017) and presynaptic homeostatic potentiation (PHP, Böhme et al., 2019; Harrell et al., 2021). In *Drosophila*, upregulated Unc-13A levels and increased numbers of Unc-13A nanomodules at presynaptic AZs in acute and chronic PHP were observed (Böhme et al., 2019). Furthermore, functional PHP is abolished in *unc-13A* null mutants and the functional dependence can be attributed to the Unc-13A N-terminus. We wanted to probe if our newly generated tool, the GFSTF-tagged *unc-13 Drosophila* strain, is suitable for further PHP analyses. To this end we measured the electrophysiological response to an acute homeostatic challenge using Philanthotoxin (PhTx) in *unc-13^GFSTF^* animals compared to control larvae (Figure 3E). We found that upon PhTx treatment *unc-13^GFSTF^* larvae showed the same increase in quantal content and thus evoked EPSC restoration as control larvae, thus, indicating that *unc-13^GFSTF^* animals can still exhibit functional PHP to the full extent. We conclude that incorporation of the GFSTF-tag into the endogenous unc-13 locus does not disrupt this form of presynaptic plasticity *in vivo*.

### Unc-13^GFSTF^ subclusters at the AZ mesoscale are reorganized during PHP

To gain further insight into the nano-arrangement of Unc-13 within the presynaptic AZ we performed two-color localization microscopy in terms of *d*STORM (Heilemann et al., 2008; van de Linde et al., 2011; Löschberger et al., 2012; Ehmann et al., 2014; Paul et al., 2015; Mrestani et al., 2021; Paul et al., 2022) using Brp^Nc82^ and a commercially available polyclonal antibody against GFP for co-localization of Brp and Unc-13^GFSTF^ (Figure 4). In previous work we established hierarchical density-based spatial clustering of applications with noise (HDBSCAN), that robustly extracts clusters in data with varying density (Campello et al., 2013; McInnes et al., 2017), as an objective way for analyzing localization data (Mrestani et al., 2021; Pauli et al., 2021; Paul et al., 2022). Here, we employed a similar approach with an extension for two-channel localization data to analyze Unc-13^GFSTF^ nanoarchitecture in the environment of the Brp scaffold (Figure 4A). First, we extracted Brp clusters in the Alexa Fluor532 channel (Figure 4Aiii, Supplementary Table 4) to remove all ‘AZ-unrelated’ Unc-13^GFSTF^ signal from our data (Figure 4iv; for detailed description see STAR Methods). We found decreased Brp cluster areas in response to PHP stimulation using Alexa Fluor532 as described before using Alexa Fluor647 (Mrestani et al., 2021, Supplementary Figure 1). The H function as derivative of Ripley’s K function was computed for the Unc-13 signal according to established algorithms (Mrestani et al., 2021) in nm steps for radii from 0 to 120 nm (Figure 4B). Next, a second HDBSCAN for extraction of individual Unc-13^GFSTF^ subclusters (SC) was applied adjusting cluster parameters until ideally matching the maximum of the H function (Figure 4C). Applying this algorithm our analysis revealed Unc-13^GFSTF^ nanoclusters with a diameter of ~26 nm which is substantially smaller than the previously described diameter of Brp SCs (Mrestani et al., 2021).

Our electrophysiological analysis revealed that Unc-13^GFSTF^ larvae can still exhibit normal PHP after PhTx treatment. Using the two-channel localization data algorithm we wanted to investigate if induction of acute PHP leads to reorganization of Unc-13^GFSTF^ clusters within the presynaptic AZ as described for other AZ components (Mrestani et al., 2021). Thus, we compared Brp and Unc-13^GFSTF^ localization data in PhTx treated preparations (phtx) and DMSO controls (ctrl, Figure 4C-E). The number of Unc-13^GFSTF^ localizations per SC as well as SC area were similar (Figure 4D) and the median number of Unc-13^GFSTF^ SCs per AZ was 11 in both groups (Figure 4E). Interestingly, we found a significantly reduced Unc-13^GFSTF^ area per AZ in phtx and the radial distance (the distance between the Brp center of mass (c.o.m.) and the Unc-13^GFSTF^ c.o.m.) was decreased (Figure 4E). Our data reveal that Unc-13^GFSTF^ forms distinct nanoclusters within the AZ in accurate co-localization with Brp which are reorganized during acute PHP.

### *d*STORM reveals formation of Unc-13^GFSTF^ superclusters at AZs

AZ circularity was comparable in Unc-13^GFSTF^ ctrl and phtx, thus, analysis of the whole dataset was plausible (Figure 5). To gain further insight into SC reorganization during PHP we focused on AZs imaged in planar view, indicated by higher circularity values (≥ 0.6, Figure 5A; compare Mrestani et al., 2021). We wondered if Unc-13^GFSTF^ SCs display a higher level of organization and performed an HDBSCAN analysis of the Unc-13^GFSTF^ SC c.o.m.s for each individual AZ (Figure 5B), thereby extracting Unc-13^GFSTF^ superclusters (SpC, Figure 5Biii). Interestingly, when comparing distance distributions of SC c.om.s we found a significant shift towards higher values in phtx compared to ctrl (Supplementary Figure 2A). Analysis of the five nearest neighbors of each individual SC c.o.m. also revealed increased distances in phtx (Figure 5C). Remarkably, whereas median and lower and 75^th^ percentiles were increased, the 90^th^ percentiles were decreased in phtx compared to ctrl (Figure 5C), indicating a narrowing of the distance distributions. This could be interpreted as a more even allocation of Unc-13^GFSTF^ SCs at the available space during PHP. Next, we asked if a higher-level organization is a feature of all AZs and Unc-13^GFSTF^ SCs. Our analysis revealed that about 50% of the Unc-13^GFSTF^ SCs per AZ were organized in about 2 SpCs that contain ~3 SCs, and that a large fraction of AZs does not display Unc-13^GFSTF^ superclustering at all (Figure 5D and Supplementary Figure 2B, C). Strikingly, the distance of SC c.o.m.s to their corresponding SpC c.o.m. was increased in phtx (Figure 5E). Taken together, we find Unc-13 SCs that are more homogenously distributed at the AZ and form enlarged SpCs correlating with increased vesicle traffic during PHP.

## DISCUSSION

We describe a MiMIC insertion of the EGFP-FlAsH-StrepII-TEV-3xFlag (GFSTF)-tag within a coding exon of the *Drosophila unc-13* gene allowing visualization of all Unc-13 isoforms expressed within the fly nervous system (Figure 1). Using SLM we demonstrate that Unc-13^GFSTF^ localizes at *Drosophila* AZs while electrophysiological characterization reveals no disturbances of spontaneous or evoked release and PHP expression (Figures 2 and 3). Furthermore, employing this newly generated tool and cluster analysis of two-channel localization data we show distinct Unc-13^GFSTF^ nanoclusters of about 26 nm diameter and ~520 nm^2^ area containing ~18 localizations at presynaptic AZs which are compacted during PHP (Figure 4).

In mammals, Munc-13 exists in nano-assemblies of 5-10 protein copies at the presynaptic AZ and is responsible for recruiting Syntaxin-1 to promote vesicle exocytosis (Sakamoto et al., 2018). It has been shown that the number of release sites equals the number of Munc13-1 clusters within the AZ and that every vesicle needs to bind to a Munc13-1 cluster to get released (Sakamoto et al., 2018). A recent study using quantitative TIRF microscopy and stepwise photobleaching underlined the functional importance of Munc13-1 clustering for vesicle docking and fusion (Li et al., 2021): a minimum of 6 Munc13-1 copies and especially the C-terminal C2C domain are necessary for efficient vesicle binding to lipid bilayers, thus, nano-clustering can be considered an inherent property of Munc13-1. Nevertheless, the exact composition and localization of Unc-13 at the AZ of an intact model organism remains to be determined.

The size of (M)unc-13 nano-assemblies is still debatable. Sakamoto et al. 2018 measured about 45 nm diameter in primary hippocampal neuronal cultures of 21 days old rats using STORM imaging. Using 2D *d*STORM imaging and the glutamatergic *Drosophila* NMJ in a newly generated GFSTF-tag MiMIC line we here describe individual Unc-13^GFSTF^ SCs of 26 nm diameter (which is well below the size of an individual synaptic vesicle in *Drosophila*, Karunanithi et al., 2002) and ~ 520 nm^2^ area (Figure 4). We used an established HDBSCAN analysis algorithm for investigation of two-channel localization data to study two AZ epitopes in close spatial relation. Brp clusters in the Alexa Fluor532 channel served as a ‘mask’ for determination of AZ extent. Remarkably, using a fluorophore with less favorable photo-physics for detection of Brp (Heilemann et al., 2008; van de Linde et al., 2011) we found similar cluster areas compared to previous Alexa Fluor647 measurements (Supplementary Table 4, compare Mrestani et al., 2021; Paul et al., 2022). The two-channel analysis allowed precise determination of the amount and extent of Unc-13^GFSTF^ SCs within the presynaptic AZ, even enabling further detailed analysis after PHP induction (see below). Distinct labeling strategies as well as the localization precision of the applied microscopic techniques might explain the different dimensions of Unc-13 clusters in our and previous work, furthermore, distinct experimental organisms and synapses have been used. *d*STORM as a variant of localization microscopy allows to count molecules (Ehmann et al., 2014; Löschberger et al., 2012). In this study we found 11-32 (25^th^-75^th^ percentile) Unc-13^GFSTF^ localizations per SC which can be translated into a certain number of Unc-13^GFSTF^ molecules. Considering the 8.1 ± 0.2 (mean ± SEM) localizations per Alexa Fluor647-labeled F(ab’)2 fragment (Ehmann et al., 2014; Löschberger et al., 2012) we assume that the 306 localizations per AZ (Supplementary Table 4) may correspond to 38 Unc-13^GFSTF^ molecules which are distributed in 11 SCs, each containing on average 3.5 Unc-13^GFSTF^ molecules. However, accounting for the usage of a polyclonal primary antibody in the present study compared to a monoclonal antibody in Ehmann et al., 2014, the true number of molecules per SC could be lower (i.e., 1 or 2 molecules). Additionally, we describe a higher-level organization that occurs in roughly 60 % of AZs, in the form of about 2 Unc-13^GFSTF^ SpCs per AZ, each containing about 3 SCs (Figure 5 and Supplementary Figure 2). Strikingly, the number of SpCs per AZ equals the previously determined number of docked vesicles (Böhme et al., 2016; Reddy-Alla et al., 2017). One might speculate that the Unc-13^GFSTF^ SpCs reported here are the mesoscale counterpart of a 6 molecule (M)unc-13 ring that, with its MUN domains aligned to 18 Synaptotagmin C2B domains, serves as a platform for SNAREpin assembly and the subsequent fusion of one synaptic vesicle (Rothman et al., 2017). The enlarged Unc-13^GFSTF^ SpC diameter from ~70 to ~74 nm (Figure 5E, Supplementary Table 4) could reflect enhanced vesicle traffic accompanied by an increased abundance of AZs captured during the process of vesicle fusion.

One also needs to consider the fact that the number of Unc-13^GFSTF^ molecules per AZ is likely to depend on the current status of the synapse, as the amounts of other essential AZ components also change in an activity-dependent manner (Kittel et al., 2006; Akbergenova et al., 2018; Rebola et al., 2019; Goel et al., 2019; Gratz et al., 2019). Previous work showed that the number of release sites *N* changes according to the applied stimuli the specific synapse faces (Atwood and Karunanithi, 2002) suggesting that the number of Unc-13^GFSTF^ clusters (equaling *N*) will also change stimulation-dependent. This study implements a newly generated and highly suitable tool, the Unc-13^GFSTF^-tag MiMIC line, to address this question of structure-function relationship in future studies. We show that Unc-13^GFSTF^ expression at the *Drosophila* NMJ is heterogeneous and varies between individual AZs (Figure 4). This finding matches the heterogeneity of synaptic release and, as the amount of other crucial AZ components like Brp seems to correlate with the amount of evoked release, creating release maps of Unc-13^GFSTF^ can be an appealing experimental approach to clarify the functional relevance of distinct nano-arrangements (Peled et al., 2014; Akbergenova et al., 2018; Newman 2022).

The Unc-13A N-terminus has been shown to be essential for expression of PHP at the *Drosophila* AZ (Böhme et al., 2019). We prove that insertion of the GFSTF-tag into the *Drosophila unc-13* gene leading to visualization of all Unc-13 isoforms does not disturb the ability of the NMJ to undergo functional PHP (Figure 3E). Specific antibodies for visualization of the Unc-13A N- and C-terminus have been published (Reddy-Alla et al., 2017; compare Figure 1C) with our Unc-13^GFSTF^ tag lying in between. Assuming localization of the N-terminal epitope close to presynaptic Ca^2+^ channels co-staining of the N-terminal antibody and Unc-13^GFSTF^ might allow visualization of the Unc-13 orientation at the presynaptic AZ. We show co-localization of all Unc-13 isoforms using the GFSTF-tag with Brp. Applying the HDBSCAN cluster analysis, approved for one-channel localization data, for the first time for two-channel localization data, our approach allows precise determination of the Unc-13^GFSTF^ nanoarrangement within the presynaptic AZ. In this study, we used PhTx for induction of acute PHP to investigate the effect on Unc-13^GFSTF^. We previously found compaction of Brp and RBP within the AZ in models of acute and chronic PHP (Mrestani et al., 2021), thus, we hypothesized that Unc-13^GFSTF^ nanoarchitecture might also change. Indeed, we found a decreased radial distance of Unc-13^GFSTF^ SCs in acute PHP accompanied by a smaller extent of the Unc-13^GFSTF^ area per entire AZ (Figure 4E). These changes reveal compaction of Unc-13 at the AZ during an acute homeostatic challenge reflecting the functional compensation of enhanced neurotransmitter release following disturbance of postsynaptic glutamate receptors (compare Mrestani et al., 2021). Future studies investigating the co-localization of Unc-13^GFSTF^ with other AZ epitopes e.g., RBP, VGCCs or RIM appear attractive to decipher the AZ nanoarchitecture. Especially in the case of less abundant proteins such as VGCCs these imaging approaches are demanding. Our labeling approach offers the possibility to co-stain different proteins using the same multi-tag GFSTF creating similar imaging conditions. Here, we used the EGFP-tag within Unc-13^GFSTF^ to visualize Unc-13 via a secondary Alexa Fluor647-labeled F(ab’)2 fragment. Further work using nanobody-staining to visualize the EGFP-tag or labeling strategies detecting the Strep-, FlaSH-tag or TEV-tag within the GFTSF-multi-tag are conceivable.

Combining the Unc-13^GFSTF^ MiMIC line with knock-out strains of crucial AZ proteins such as RIM or RBP to elucidate their effects on the amount or expression pattern of Unc-13 appears promising. Due to the interaction of Munc-13, Rab3 and RIM in mammals the RIM knock-out is likely to affect the Munc-13 nanotopology, however, this is unclear in *Drosophila* as formation of the tripartite complex is lacking. The RBP knock-out should affect Unc-13^GFSTF^ expression in the fly (Petzoldt et al., 2020) which can be addressed using the here described MiMIC line. This genetic tool will be useful for further studies analyzing AZ nanoarchitecture and structure-function relationships at the *Drosophila* NMJ.

## ABBREVIATIONS

AZ: active zone
Brp: Bruchpilot
Ca^2+^: Calcium
CaM: Calmodulin
DMSO: dimethyl sulfoxide
CRISPR: clustered regularly interspaced short palindromic repeats
*d*STORM: *direct* stochastic optical reconstruction microscopy
eEPSC: evoked excitatory postsynaptic current
GFP: green fluorescent protein
sfGFP: superfolder GFP
EGFP: enhanced GFP
GFSTF: EGFP-FlAsH-StrepII-TEV-3xFlag
GluRIIA: glutamate receptor IIA subunit
HDBSCAN: hierarchical density-based spatial clustering of applications with noise
mEPSC: miniature excitatory postsynaptic current
MiMIC: *Minos-mediated* integration cassette
NMJ: neuromuscular junction
PHP: presynaptic homeostatic potentiation
PhTx: Philanthotoxin 433 tris (trifluoroacetate) salt
RIM: Rab3-interacting molecule
RMCE: recombinase-mediated cassette exchange
SC: subcluster
SIM: structured illumination microscopy
SLM: super-resolution light microscopy
SNARE: soluble N-ethylmaleimide-sensitive-factor attachment receptor
SpC: supercluster
SV: synaptic vesicle
TEVC: two-electrode-voltage clamp
VGCC: voltage-gated calcium channels

## AUTHOR CONTRIBUTIONS

SD, AM, MH and MMP designed experiments. SD, AM, FG, MP, FK and MMP performed experiments. SD, AM, FG, MP, PK, MS and MMP analyzed the data. SD, AM and MMP wrote the manuscript with help of all co-authors. MH and MMP coordinated the study and provided funding.

## ACKNOWLEDGEMENTS

This work was supported by grants from the German Research Foundation (DFG) FOR 3004 SYNABS P1 to MH, the IZKF Würzburg to MH (N229) and MMP (Z-3/69) and the University of Leipzig Clinician Scientist Program to AM. The authors thank Tobias Langenhan for scientific discussions and Maria Oppmann and Frauke Köhler for technical assistance.

## SUPPLEMENTARY MATERIAL

**Supplementary Figure 1.**
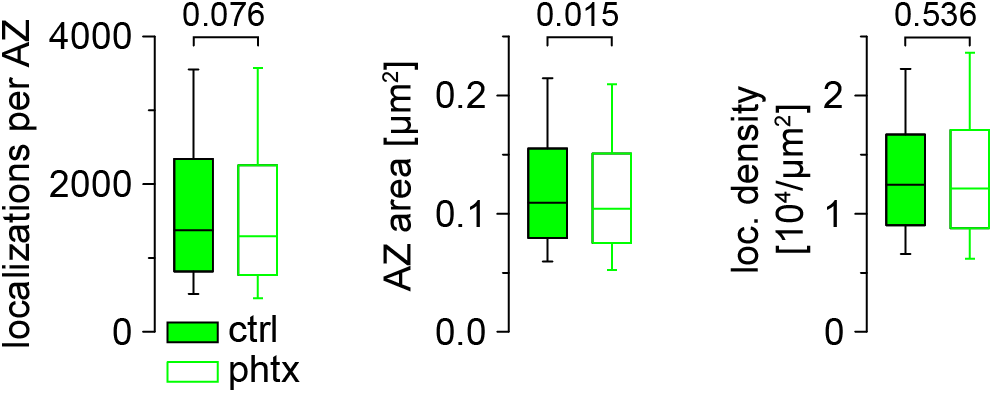
Active zone compaction using the dye Alexa Fluor532. Number of Brp localizations per AZ, AZ area and Brp localization density for ctrl (filled boxes, n = 1,462 AZs from 22 NMJs and 9 animals) and phtx (open boxes, n = 1,521 AZs from 23 NMJs and 9 animals) shown as shown as box plots (horizontal lines show median, boxes boundaries 25^th^ and 75^th^ percentiles and whiskers 10^th^ and 90^th^ percentiles) imaged in *unc-13^GFSTF^* type Ib boutons stained with Brp^Nc82^ labelled with Alexa Fluor532 conjugated IgGs (green channel from data in Figure 4).

**Supplementary Figure 2.**
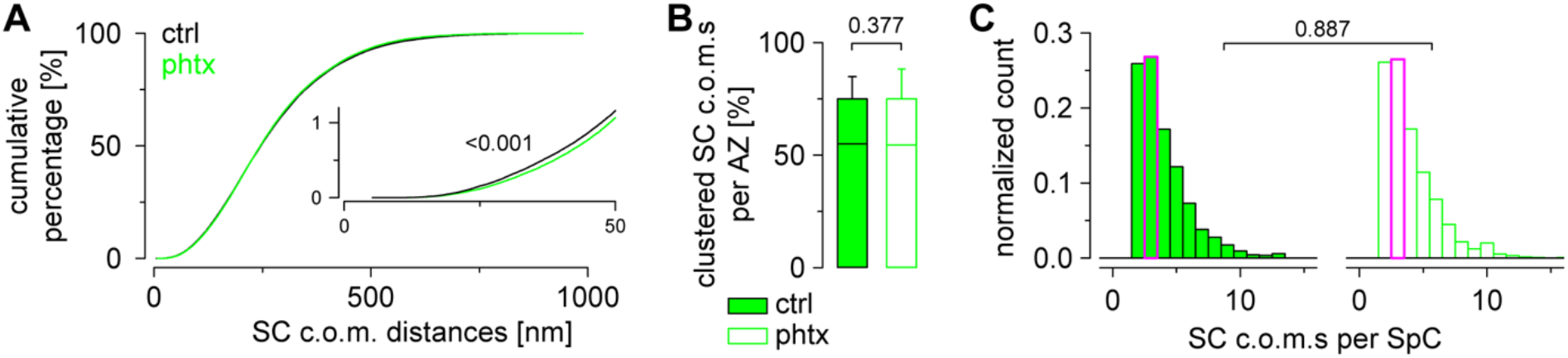
Extended Unc-13 supercluster analysis. (A) Cumulative plots show distance distributions between Unc-13^GFSTF^ SC c.o.m.s at circular AZs for ctrl (n = 162,170 distances from 10,461 SCs and 876 AZs) and phtx (n = 169,874 distances from 10,988 SCs and 927 AZs). Inset shows x and y axes segments 0 and 50 nm and 0 and 1.2 %, respectively. (B) Percentage of SC c.o.m.s that are organized in SpCs per AZ for ctrl (n = 876 AZs from 22 NMJs and 9 animals) and phtx (n = 927 AZs from 23 NMJs and 9 animals). (C) Number of SC c.o.m.s per SpC for both groups at AZs with at least one SpC (n = 1,417 and 1,497 SpCs for ctrl and phtx, respectively).

**Supplementary Table 1.**
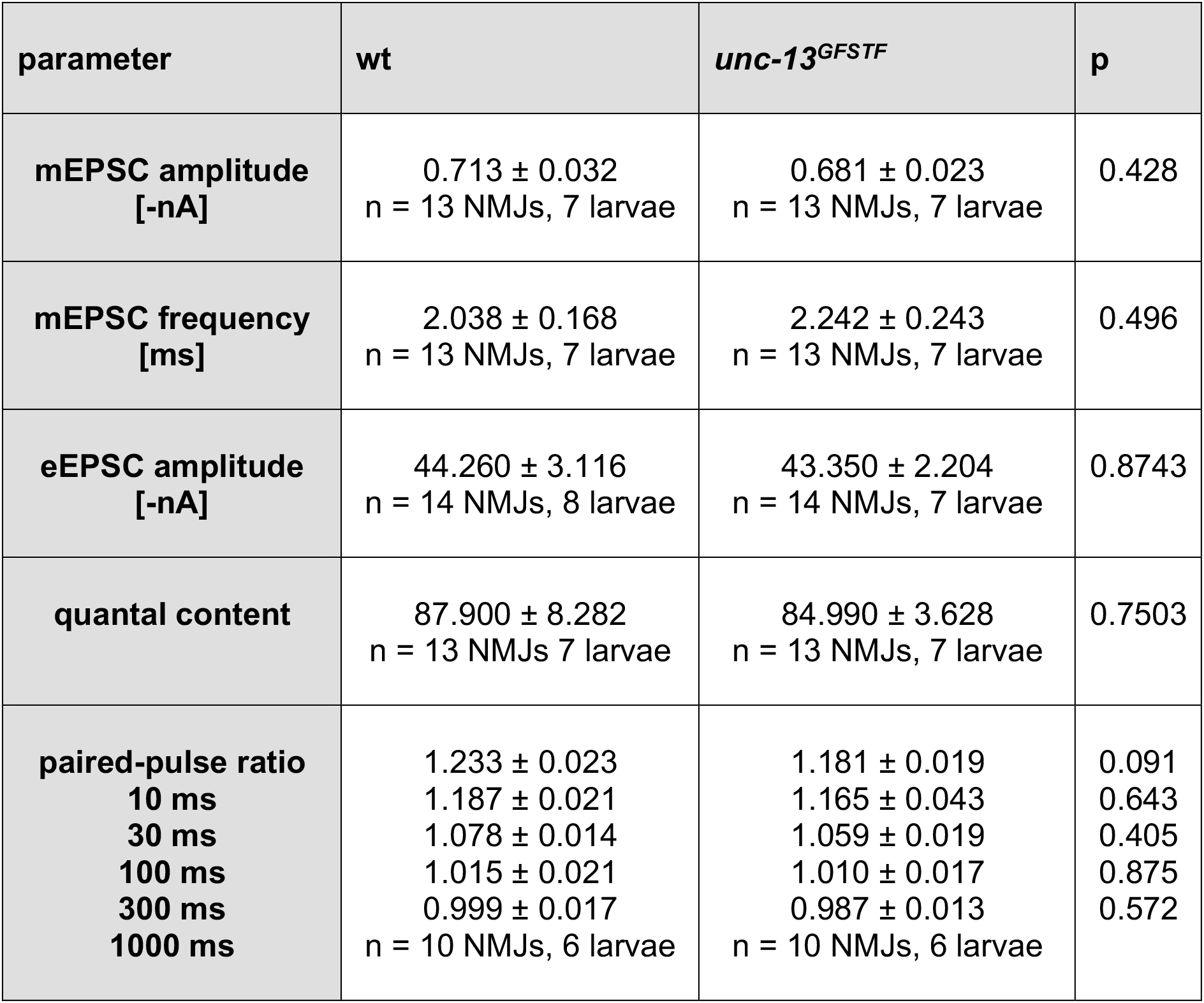
Electrophysiological analysis of spontaneous and evoked synaptic transmission in *unc-13^GFSTF^* and wildtype larvae. Related to Figure 3A-D. Numerical values are given as mean ± SEM for each genotype. p-values and sample sizes for the number of NMJs and the number of animals used for analysis are indicated. Data were normally distributed and unpaired t-tests were used to evaluate statistical significance.

**Supplementary Table 2.** Electrophysiological analysis of acute presynaptic homeostasis in *unc-13^GFSTF^* and wildtype animals. Related to Figure 3E. Numerical values are given as mean ± SEM for each group. p-values and sample sizes for the number of NMJs and the number of animals used for analysis are indicated. For information of statistical analysis see Supplementary Table 3.

| parameter | wt (DMSO) | wt (PhTx) | *unc-13<sup>GFSTF</sup>* (DMSO) | *unc-13<sup>GFSTF</sup>* (PhTx) |
| --- | --- | --- | --- | --- |
| **eEPSC amplitude [-nA]** | 37.83 ± 2.45<br>n = 10 NMJs,<br>7 larvae | 44.20 ± 2.31<br>n = 10 NMJs,<br>5 larvae | 41.99 ± 2.26<br>n = 13 NMJs,<br>6 larvae | 47.14 ± 2.88<br>n = 14<br>NMJs, 7<br>larvae |
| **mEPSC amplitude [-nA]** | 0.647 ± 0.065<br>n = 10 NMJs,<br>7 larvae | 0.413 ± 0.020<br>n = 10 NMJs,<br>5 larvae | 0.716 ± 0.033<br>n = 13 NMJs,<br>6 larvae | 0.393 ±<br>0.017, n =<br>14 NMJs, 7<br>larvae |
| **quantal content** | 84.89 ± 9.82<br>n = 10 NMJs,<br>7 larvae | 144.90 ± 9.03<br>n = 10 NMJs,<br>5 larvae | 80.25 ± 5.47<br>n = 13 NMJs,<br>6 larvae | 165.50 ±<br>14.02, n =<br>14 NMJs, 7<br>larvae |

**Supplementary Table 3.** Statistical comparison of acute presynaptic homeostasis in *unc-13^GFSTF^* and wildtype animals. Related to Figure 3E. p-values revealed by parametric t-tests (eEPSC amplitude) or by Mann-Whitney Rank Sum tests for non-parametric data (mEPSC amplitude and quantal content) are given for comparisons between both genotypes or between measurements in DMSO and PhTx within an individual genotype.

| genotype | eEPSC<br>amplitude | mEPSC<br>amplitude | quantal content |
| --- | --- | --- | --- |
| wt (DMSO)<br>vs. wt (PhTx)<br>vs. *unc-13<sup>GFSTF</sup>* (DMSO) | 0.075<br>0.229 | 0.002<br>0.257 | 0.001<br>0.784 |
| wt (PhTx)<br>vs. *unc-13<sup>GFSTF</sup>* (PhTx) | 0.462 | 0.472 | 0.472 |
| *unc-13<sup>GFSTF</sup>* (DMSO)<br>vs. *unc-13<sup>GFSTF</sup>* (PhTx) | 0.176 | < 0.0001 | < 0.0001 |

**Supplementary Table 4.**
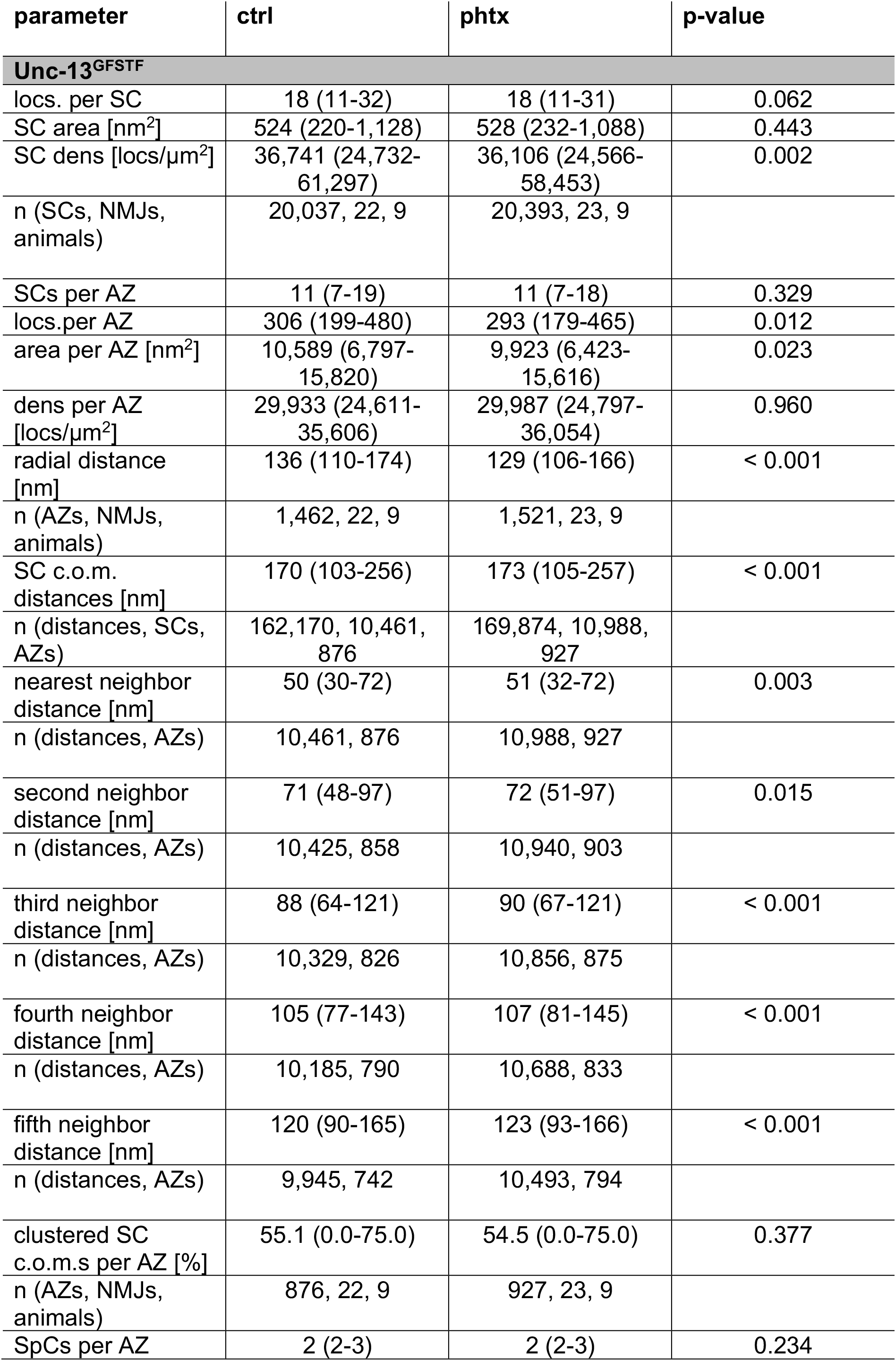
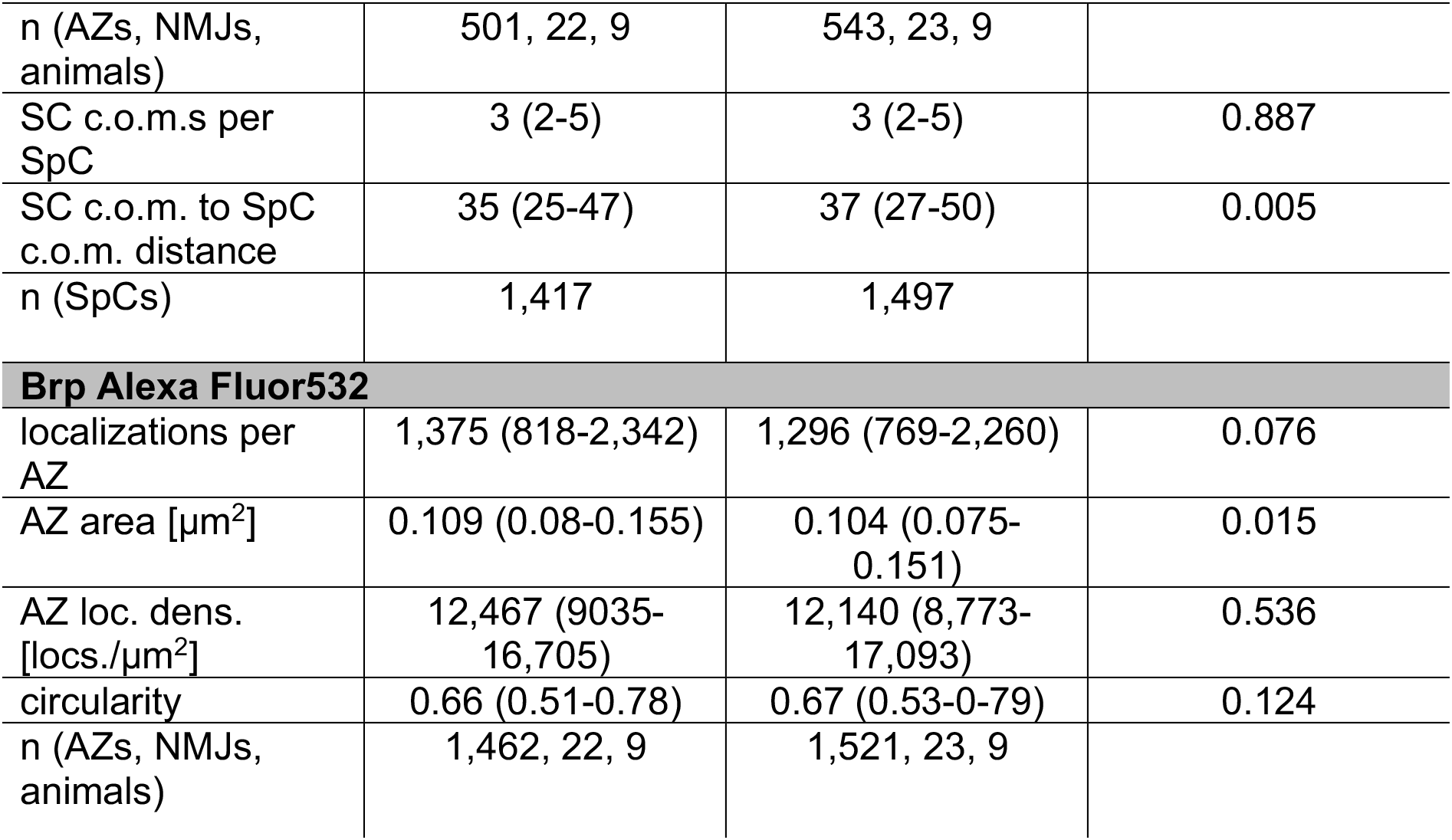
*d*STORM analysis of Unc-13^GFSTF^ subclusters using Alexa Fluor647 and Brp clusters using Alexa Fluor532. Related to Figures 4 and 5 and Supplementary Figure 2. Non-parametric data, reported as median (25^th^-75^th^ percentile).

